# Neonatal enteropathogenic *Escherichia coli* infection disrupts microbiota-gut-brain axis signaling

**DOI:** 10.1101/2020.09.08.288803

**Authors:** Carly Hennessey, Mariana Barboza, Ingrid Brust-Mascher, Trina A Knotts, Jessica A. Sladek, Matteo M Pusceddu, Patricia Stokes, Gonzalo Rabasa, Mackenzie Honeycutt, Olivia Walsh, Colin Reardon, Mélanie G. Gareau

## Abstract

**Background:** Diarrheal diseases are a leading cause of death in children under age five worldwide. Repeated early life exposures to diarrheal pathogens can result in co-morbidities including stunted growth and cognitive deficits suggesting an impairment in the microbiota-gut-brain (MGB) axis.

**Methods:** Neonatal C57BL/6 mice were infected with EPEC (strain e2348/69; ΔescV [T3SS mutant]), or vehicle (LB broth) via orogastric gavage (10^5^ CFU) at post-natal day (P7). Behavior (novel object recognition [NOR] task, light/dark [L/D] box, and open field test [OFT]), intestinal physiology (Ussing chambers), and the microbiota (16S Illumina sequencing) were assessed in adulthood (6-8 weeks).

**Results:** Neonatal infection of mice with EPEC impaired recognition memory (NOR task), coupled with increased neurogenesis (Ki67 and doublecortin immunostaining) and neuroinflammation (increased microglia activation [Iba1]) in adulthood. Intestinal pathophysiology was characterized by increased secretory state (short circuit current; Isc) and permeability (conductance; FITC-dextran flux) in the ileum and colon of neonatally EPEC-infected mice, along with increased expression of pro-inflammatory cytokines (*Tnfα, Il12, Il6*) and pattern recognition receptors (*Nlr, Tlr*). Finally, neonatal EPEC infection caused significant dysbiosis of the gut microbiota, including decreased Firmicutes, in adulthood.

**Conclusions:** Together these findings demonstrate that infection in early life can significantly impair the MGB axis in adulthood.

## INTRODUCTION

Despite advances in sanitation and access to clean drinking water, enteric infections remain a global public health issue, with diarrheal diseases a leading cause of death in children under age five worldwide (1). While treatment of diarrhea with oral rehydration solution has significantly decreased childhood mortality, repeated exposures to diarrheal pathogens have been correlated with impaired growth and cognitive deficits into adulthood (2-4). Malnutrition caused by enteric bacterial pathogens may support ongoing dysbiosis, resulting in gastrointestinal (GI) barrier dysfunction and systemic translocation of bacterial products (3, 5). This translocation may drive systemic inflammation, leading to neuroinflammation, which could contribute to cognitive deficits. Despite this association, the underlying cause of enteric infection-induced cognitive deficits is unknown.

Early neonatal life is a critical time for neural development (6), differentiation and maturation of the GI tract (7), and colonization of the gut microbiota (8), together comprising the microbiota-gut-brain (MGB) axis. Signaling of the MGB axis occurs via hormones, neurotransmitters, and immune cells, although the precise pathways involved remain to be fully elucidated (9, 10). Dysbiosis, induced following either antibiotic administration (11) or infection with a bacterial pathogen (12) can result in long-lasting impairments in MGB axis signaling, including behavioral deficits and GI pathophysiology. For example, in adult mice, infection with the murine bacterial pathogen, *Citrobacter rodentium*, causes acute anxiety-like behavior (13) and stress-induced cognitive deficits (12), with exposure to social stress increasing severity of infection (14). These studies highlight the association between infection, dysbiosis, inflammation, and behavioral impairments, which may be further enhanced following initiation during early neonatal life.

Here, we sought to identify the effects of neonatal bacterial enteric infection on the MGB axis by infecting mouse pups with enteropathogenic *Escherichia coli* (EPEC). While EPEC does not typically colonize adult mice without antibiotic pre-treatment (15), it can effectively colonize both ileum and colon in neonatal mice (16). Epithelial cell attachment and EPEC microcolony formation depend on the presence of a functional bundle forming pili and type III secretion system (T3SS) (16). Using a similar neonatal EPEC infection model, we identified recognition memory deficits, GI pathophysiology, and dysbiosis in adult mice. These MGB axis defects were coupled with increased hippocampal neurogenesis and neuroinflammation, identifying a potential novel pathway through which a neonatal enteric bacterial infection can cause cognitive deficits in adulthood.

## METHODS

### Mice

Mice (male and female C57BL/6J; originally from Jackson Laboratories) were bred in-house. Mice were housed in cages lined with chip bedding, exposed to 12h light/dark cycle, and had access to food and water *ad libitum*. Mice were euthanized by CO_2_ followed by cervical dislocation. All procedures and protocols were approved by UC Davis (IACUC #20072).

### Study Design

Neonatal mice were challenged on P7 with 50µl of EPEC (O127:H6, strain E2348/69 or T3SS mutant [ΔescV; kindly provided by Dr. Andreas Baumler]; 10^5^ colony forming units [CFU]) or vehicle (Luria-Burtani [LB] broth) via orogastric gavage (**Fig. 1A**). Adult mice (6-8 weeks) were used for three sets of experiments:

**Figure 1:**
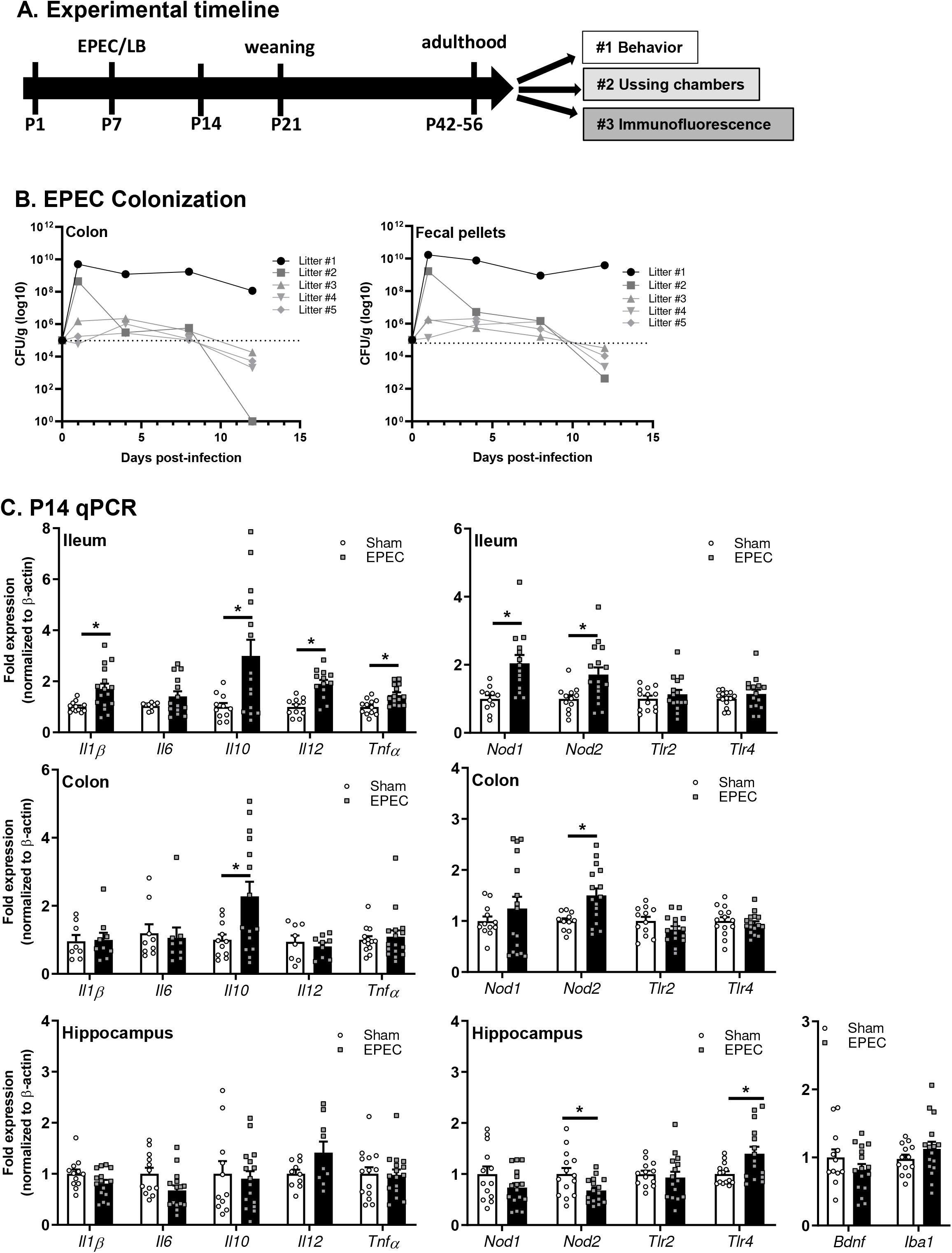
Study design and EPEC colonization. (**A**) Study design. **(B)** Colonies from fecal samples grown on MacConkey agar plates 1, 4, 8, and 12 days post infection were counted to quantify EPEC colonization. Data represents n=1-2 pups per time point from 5 separate litters and dotted line represents infectious dose (10^5^ CFU). (**C**) Relative mRNA expression in ileum, colon, and hippocampus at P14. (N=12-16; *p<0.05 by Student’s T test (± Welch’s correction), or Mann-Whitney where appropriate).

1. *Behavioral testing*. Mice were acclimated to the testing room overnight. Behavior was assessed during the light cycle (9am - 6pm) from least to most stressful: light-dark (L/D) box, open field test (OFT), and novel object recognition (NOR) task. This strategy prevents carry-over effect between tests (17). Tissues (hippocampus, distal ileum and proximal colon) were collected for qPCR within 30 minutes post-behavioral testing.
2. *GI pathophysiology*. Distal ileum and proximal colon were collected for Ussing chamber studies.
3. *Neuroinflammation and neurogenesis*. Mice were anaesthetized with isoflurane (5%), perfused with 4% paraformalehyde (PFA), and brains collected.

### EPEC Colonization

To monitor EPEC infection, colonization was assessed at 1, 4, 8, and 12 days post infection (DPI) in five separate litters. Proximal colon samples (tissue ± fecal pellets) were weighed, homogenized, and serially diluted (10^−1^ to 10^−5^) for plating on MacConkey agar and incubated at 37°C overnight. Data is presented as CFU/g tissue.

### Behavior Tests

#### L/D Box

Anxiety-like behavior was assessed using the L/D box test (17-19). Briefly, mice were placed in a novel arena (40 × 20 cm) divided into a dark compartment (1/3) and a light compartment (2/3) for 10 minutes. Automated tracking software (Ethovision, Noldus) was used to measure the time spent in the light box and count the number of transitions between the light and dark compartments.

#### OFT

General locomotor activity was assessed using the OFT (17). Briefly, mice were placed in a novel arena (30 ⨯ 30 cm) and allowed to explore for 10 minutes. Locomotor activity was assessed by measuring total distance traveled (m). Time spent in the center zone (12⨯ 12 cm), and number of transitions into the center zone served as secondary measures of anxiety-like behavior. Automated tracking software (Ethovision, Noldus) was used to measure parameters.

#### NOR Task

Recognition memory was assessed using the NOR task (17, 19, 20). Mice were placed in an arena (30 × 30 cm) and allowed to acclimate. Behavior during the acclimation period was recorded and analyzed as part of the OFT (described above). The NOR task consisted of two phases: training and testing. In the training phase, mice explored two identical objects for 5 minutes, followed by 45 minutes of rest in their home cage. During the testing phase, a familiar object was replaced with a novel object and behavior was recorded for 5 minutes. The number of interactions with each object in the training and testing phase was scored manually (JS and CH); with an exploration ratio calculated as the number of interactions with the new object divided by total number of interactions. Only mice that passed recognition criteria (no preference for either object [40-60% recognition] during the training phase and at least 10 interactions with both objects in either phase) were included.

#### Ussing chambers

Distal ileum and proximal colon were excised, cut along the mesenteric border, and mounted onto cassettes (Physiologic Instruments, San Diego, CA) for Ussing chamber studies as described previously (17, 21). After 15 minutes of equilibration, 4KDa FITC-labeled dextran (88mg/ml; FD4, Sigma) was added to the mucosal chamber to measure macromolecular permeability over time. Baseline active ion transport was measured by short circuit current (Isc, μA/cm^2^) and tight junction permeability was measured by conductance (G, mS/cm^2^) using Acquire and Analyze software (Physiologic Instruments). Serosal samples were collected every 30 minutes for 2 hours with FITC dextran flux (ng/ml/cm^2^/hr) determined by plate reader (Biotek Synergy H1).

### qPCR

GI (distal ileum and proximal colon) and brain (hippocampus, pre-frontal cortex [PFC], cerebellum) were collected and stored in Trizol (Invitrogen, Carlsbad, CA) at −80°C. RNA was extracted according to manufacturer’s protocol (Invitrogen) and cDNA prepared using iScript cDNA transcriptase kit (BioRad). Gene expression was determined by qPCR (Applied Biosystems QuantStudio 6 Flex) with SYBR Green master mix using primers from Primerbank (22) (**Supplemental Table 1**). Data was analyzed using 2^−ΔΔCt^ method and presented as fold change relative to β-actin.

### Immunofluorescence

Perfused brains were post-fixed in 4% PFA (48h), and cryoprotected in 30% sucrose + 0.1% sodium azide in PBS (48h). Subsequently, brains were embedded in optimal cutting temperature (OCT) medium (Fisher Scientific), and stored at −20°C. Coronal sections (40μm) were cut using a cryostat (Leica, Germany).

#### Neurogenesis

Neurogenesis was assessed by quantifying the number of proliferating cells (Ki67^+^) and immature neurons (doublecortin [DCX]^+^) in the dentate gyrus (DG) of the hippocampus (17). Coronal sections were co-stained with primary antibodies (rabbit anti-Ki67 [LS-C141898, Lifespan Biosciences, Seattle, WA] and guinea pig anti-DCX [AB2253, Millipore, Burlington, MA]), followed by secondary antibodies (Ki67: Alexa 647 goat anti-rabbit [ab150155, Abcam, Cambridge, UK]; DCX: Alexa 555 goat anti-guinea pig [A21435, Invitrogen]) and nuclei were stained with DAPI. Confocal images were taken with a 20X objective (Leica, SP8), 1.04 µm z-step size, with the number of Ki67^+^ and DCX^+^ cells per DG volume quantified using Imaris (8.2.1) (23) and presented as number of positive cells per μm^3^.

#### Neuroinflammation

Microglia morphology was used to assess hippocampal neuroinflammation. Coronal brain sections were stained with rabbit anti-Iba-1 (019-19741, Wako), followed by secondary antibody (Alexa Fluor 546 donkey anti-rabbit [Invitrogen]), and DAPI for nuclear staining. Confocal images were taken with a 63X objective and 0.3 μm z-steps (Leica SP8). 3-D reconstructions of microglia was performed using a modified version of the open source 3DMorph Matlab script (24) and Imaris software for cell quantification, and characterization of processes (length, numbers, branch points) in both the DG and the CA1 regions (21).

### Microbiome analysis

Fecal samples were collected at euthanasia, frozen at −80°C, and sent to Molecular Research LLC (MR DNA) for processing and sequencing. DNA was extracted using the PowerSoil fecal extraction kit (Qiagen), with library preparation and sequencing performed using primer pair 515F and 806R in 300 bp paired end run on an Illumina MiSeq (Illumina, San Diego). Sequencing data was analyzed using QIIME2 version 2019.7.0 (25) with demultiplexed paired end sequences with barcodes and adapters removed. DADA2 was used for quality filtering and feature (OTU) prediction (26). Based on sequence quality, forward reads were truncated to 220 nts and the reverse reads were truncated to 230 nts. MAFFT was used to align representative sequences (27), which were organized into a phylogenetic tree using FastTree 2 (28). Taxonomic classification using OTUs/features was performed using a pre-trained Naïve Bayes taxonomy classifier: Silva132_99%OTUs (29), to generate taxonomic counts and percentage (relative frequency) tables. Diversity analyses were run on the resulting OTU/feature.biom tables to provide both phylogenetic and non-phylogenetic metrics of alpha- and beta-diversity (30). Raw sequences have been uploaded to the NCBI sequence read archive under accession number PRJNA478451.

### Statistical analysis

Results are presented as averages ± SEM. Student’s T-test (with Welch’s correction for unequal variances) and Mann-Whitney test (non-parametric data) were performed where appropriate using Prism V.8 (GraphPad, San Diego, CA). Kruskal-Wallis pairwise comparison was used for microbiome analysis. P<0.05 was considered to be statistically significant.

## RESULTS

### EPEC colonizes neonatal mice and leads to GI inflammation

EPEC colonization was assessed over time until weaning. EPEC effectively colonized mice, as demonstrated by increased CFU/g tissue, reaching 10^6^-10^8^ CFU/g feces by 1-4 DPI and clearance starting at 12 DPI **(Fig. 1B)**. At P14, 7 DPI, expression of *Il1β, Il10, Il12*, and *Tnfα* was significantly increased in the ileum of EPEC-infected vs. sham-infected mice (**Fig. 1C**). In the proximal colon, immune activation was considerably limited in EPEC-infected mice with only increased Il10 expression (**Fig. 1C**), while no evidence of immune activation was seen in the hippocampus. Infection also significantly increased the expression of the pattern recognition receptors (PRR) *Nod1* and *Nod2* in the ileum and *Nod2* in the colon, whereas in the hippocampus, EPEC infection reduced *Nod2* expression and increased *Tlr4*. Inflammation was dampened by P21, with only *Il1β* in the ileum and *Il6, Il10*, and *Tnfα* elevated in the colon as bacterial burden decreased (**Suppl Fig. 1**). In the hippocampus, elevated Il1*β* and *Il6* were seen along with increased expression of the microglia marker *Iba1* (**Suppl Fig. 1**), suggesting ongoing mild immune activation at P21 in both the gut and the brain.

### Neonatal EPEC infection causes cognitive deficits in adulthood

To study the impact of neonatal EPEC infection on the MGB axis in adulthood, behavioral testing was performed. Recognition memory (NOR task), was impaired in EPEC-infected mice versus sham-infected controls as indicated by a decreased exploration ratio (**Fig. 2A**). This effect was dependent on a T3SS, as recognition memory was unaffected in mice infected with the ΔescV mutant strain (sham: 58.26±1.10% vs ΔescV: 56.33±1.30%; N=20-23). In contrast, EPEC infection did not induce anxiety-like behavior (L/D box), with time spent in the light and transitions between the light and dark boxes similar between sham and EPEC groups (**Fig. 2B**). Locomotor activity (OFT) was also normal, with total distance travelled, frequency of transitions, and time spent in the center of the arena the same in EPEC-infected mice as sham-treated controls (**Fig. 2C**). These findings suggest that neonatal EPEC infection causes recognition memory deficits in adulthood in a T3SS-dependent manner, without inducing anxiety-like behavior or reducing exploratory behaviors.

**Figure 2.**
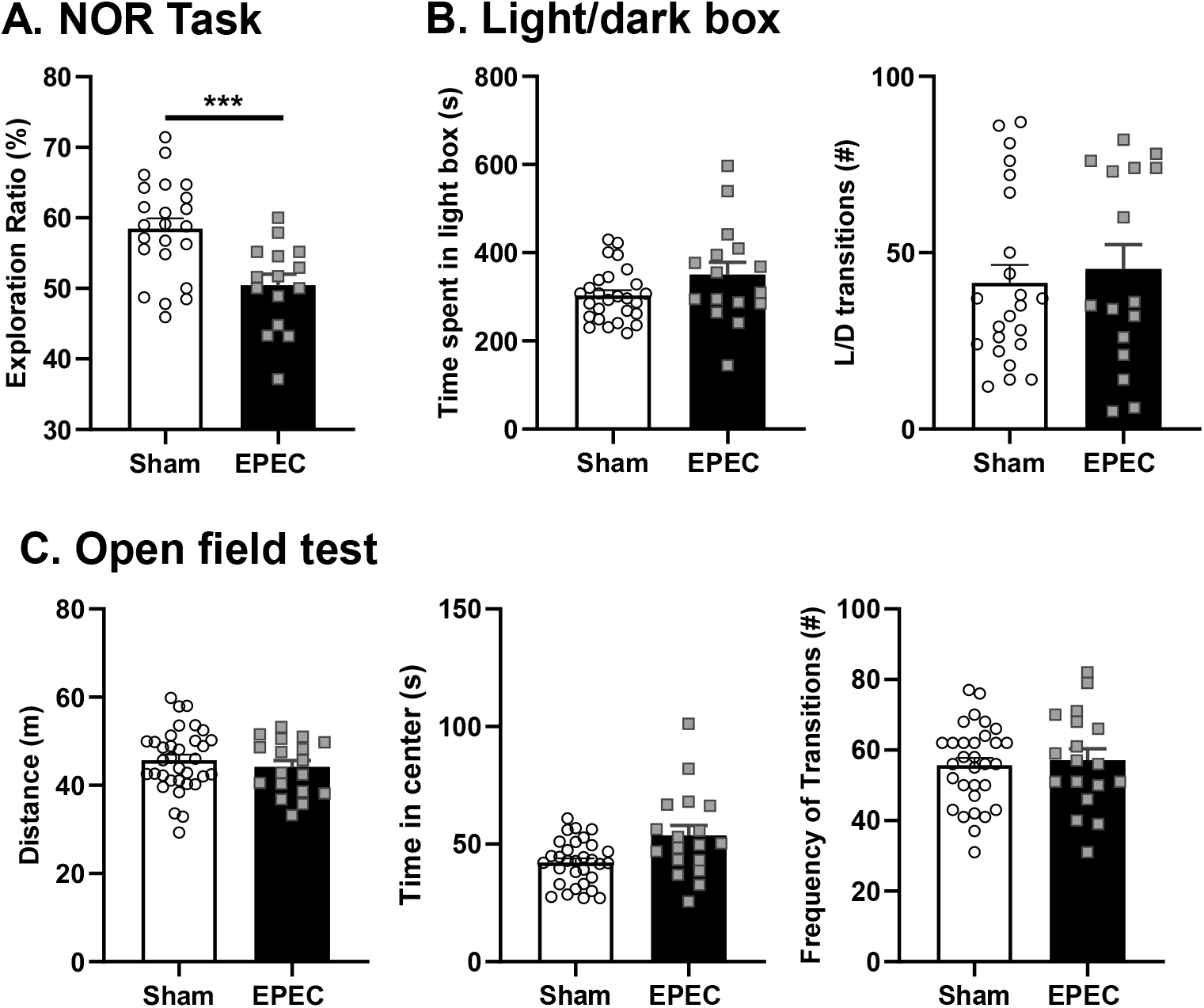
Neonatal EPEC infection induces cognitive deficits in adulthood. (**A**) Recognition memory was assessed using the novel object recognition (NOR) task. (**B**) Anxiety-like behavior was assessed using the light/dark (L/D) box to measure time spent in the light box and transitions between the light and dark box. (**C**) General behavior and locomotor activity were determined using the open field test (OFT), including distance traveled, frequency of transitions to the inner zone, and time spent in the inner zone. (N=15-34, ***p<0.001 by Student’s T-test.

### Neonatal EPEC infection leads to increased neurogenesis in adult hippocampus

As the primary site for adult neurogenesis, the DG region of the hippocampus plays a critical role in hippocampal plasticity (31) as well as learning and memory (32). Immature neurons (DCX^+^/μm^3^; *p<0.05) and proliferating cells (Ki67^+^/μm^3^; *p<0.05) were significantly increased in the DG of EPEC-infected mice versus sham-infected controls (**Fig. 3A-B**). Increased neurogenesis in EPEC-infected mice was associated with increased expression of brain derived neurotrophic factor (BDNF), critical for supporting neuronal growth and neurogenesis (32) (**Fig. 3C**). Taken together, these findings suggest that neonatal EPEC infection increases hippocampal neurogenesis in adulthood.

**Figure 3.**
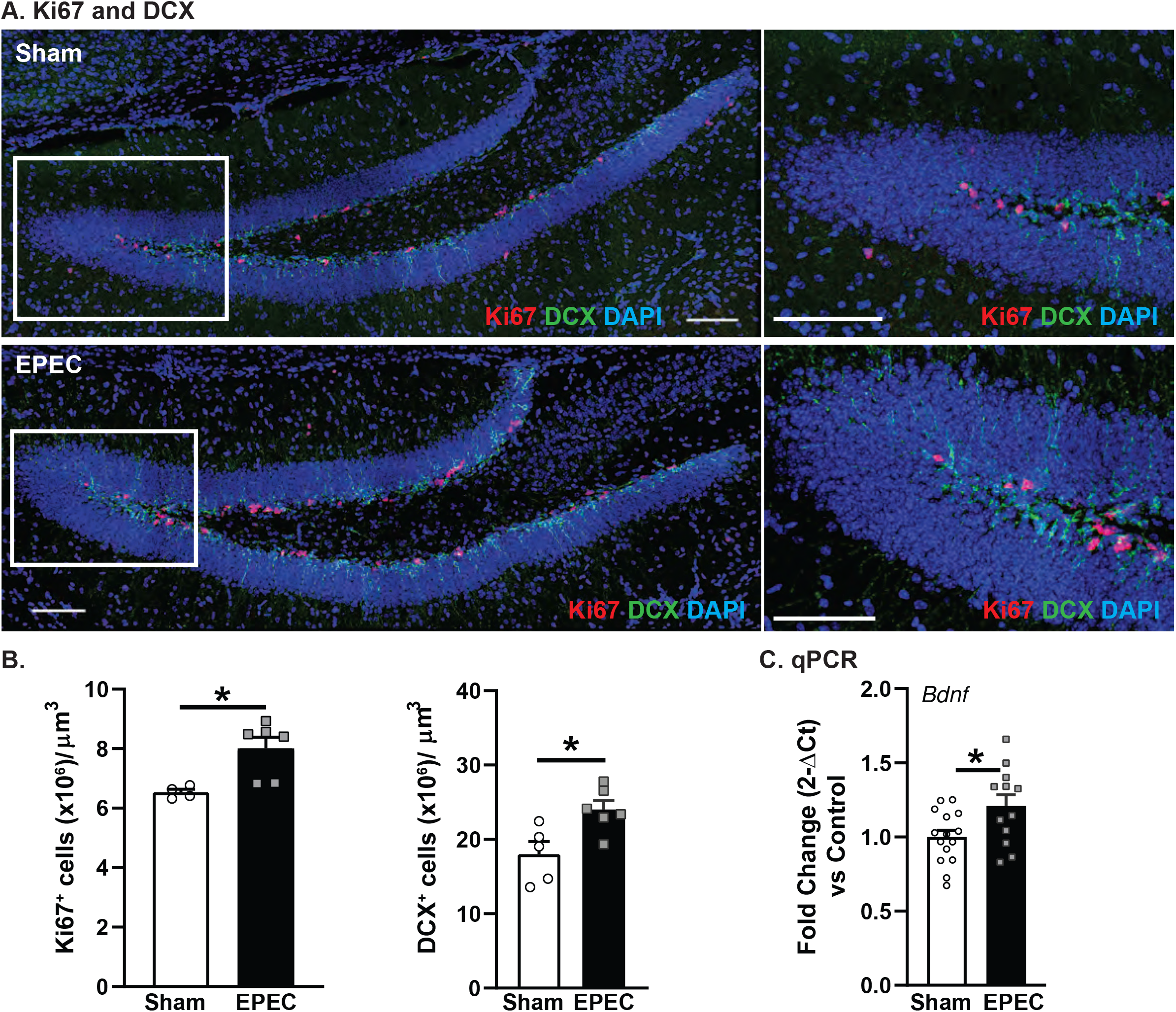
Neonatal EPEC infection impairs hippocampal neurogenesis in adulthood. (**A**) Confocal imaging for markers of neurogenesis, including Ki67 (cell proliferation) and doublecortin (DCX, immature neurons) by immunofluorescence (scale bars = 100 μm). (**B**) Quantification of Ki67^+^ and DCX^+^ cells per cubic micron in the dentate gyrus (DG; N=5; *p<0.05; Student’s T-test). (**C**) Relative mRNA expression of *Bdnf* in the hippocampus (N=16; *p<0.05; Student’s T-test).

### Neonatal EPEC infection causes neuroinflammation and microglia activation in adulthood

Since neonatal EPEC infection reduced recognition memory and increased hippocampal neurogenesis in adulthood, neuroinflammation was assessed by qPCR. *Although hippocampal expression of Il10* and *Il22 were both increased, no change was observed in Il1β, Il6, or TNFα in adult mice neonatally infected with EPEC*. Increased expression of several PRRs (*Nod1, Nod2, and Tlr2*) was also observed, along with increased expression of the microglia marker Iba1 (**Fig. 4A**) in adult EPEC mice, suggesting immune activation. This was specific to the hippocampus, with the cerebellum and PFC region showing little evidence of immune activation (**Suppl Fig. 2**) Microglia are resident immune cells in the brain, with their activation regulated in part by the gut microbiota (33). Upon activation, microglia retract their processes becoming more amoeboid, changing their morphology (33). Hippocampal microglia in EPEC-infected mice had a more activated phenotype, characterized by significantly decreased process lengths, decreased number of branch points, and reduced number of terminal points in the DG and CA1 regions, coupled with increased cell numbers (**Fig. 4B**). Taken together, these findings support activation of microglia in the adult hippocampus following neonatal EPEC infection.

**Figure 4.**
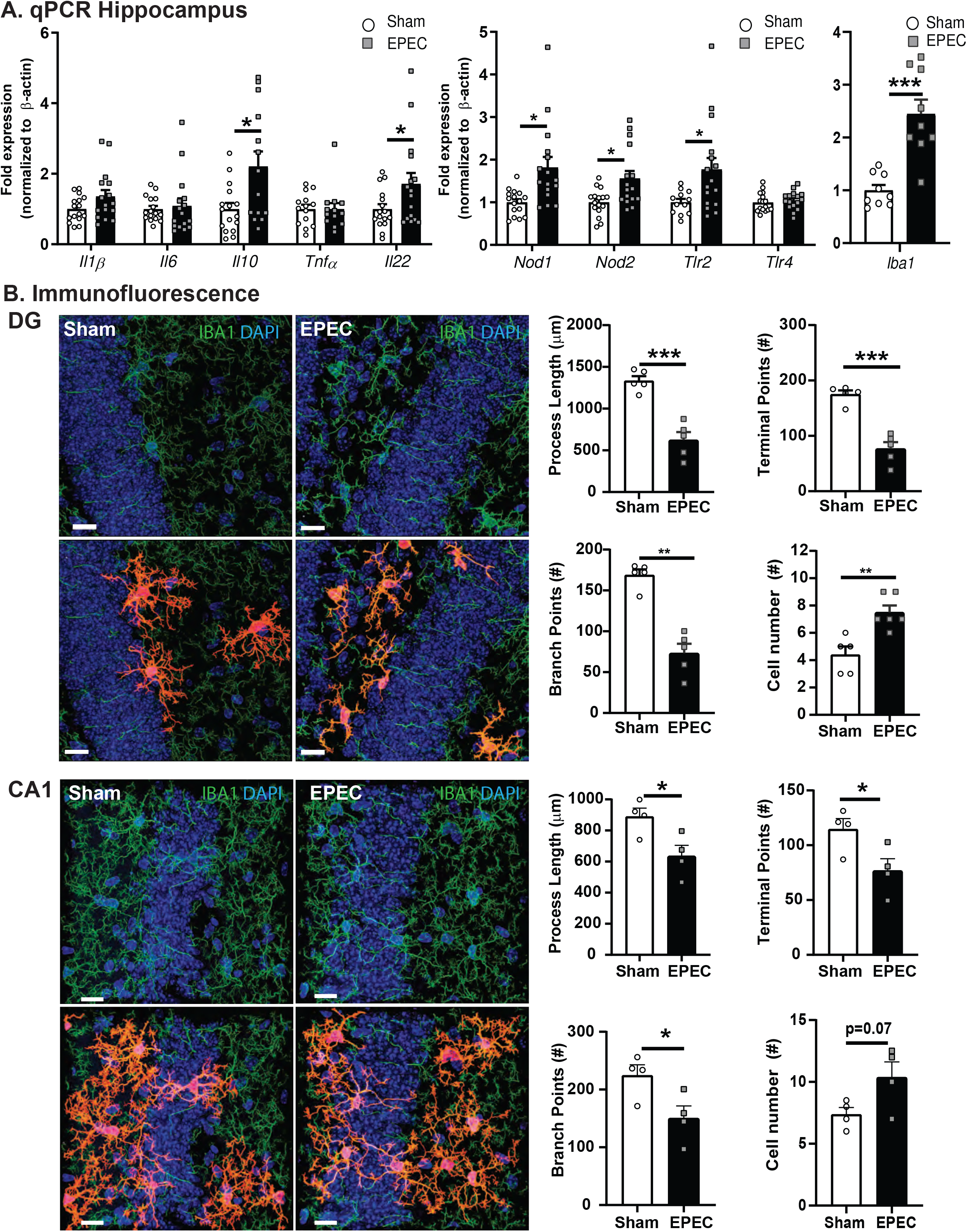
Neonatal EPEC infection leads to neuroinflammation. (**A**) Relative mRNA expression in the hippocampus (N=12-16; *p<0.05, **p<0.01; Student’s T-test, Mann-Whitney and Welch’s correction where appropriate). (**B**) Confocal imaging and morphological characterization of microglia in the dentate gyrus (DG) and CA1 region of the hippocampus (N=5-6; **p<0.01, ***p<0.001; Student’s T-test, scale bars =15 μm).

### Neonatal EPEC infection results in intestinal pathophysiology in adulthood

Diarrheal disease is caused by an imbalance between secretory and absorptive functions of the intestinal epithelium, often coupled with impaired permeability (34). Neonatal EPEC infection resulted in increased baseline ion transport (Isc) in the ileum and colon compared to sham-infected controls **(Fig. 5A**). Increased permeability was also demonstrated in neonatally EPEC-infected mice by elevated G **(Fig. 5B**) and FITC-dextran flux (**Fig. 5C**) in both ileum and colon, indicating increased tight junction and macromolecular permeability, respectively. Neonatal EPEC infection also resulted in increased expression of pro-inflammatory cytokines (*Il6, Il22, Tnfα)*, PRRs (*Nod1, Nod2, Tlr2, Tlr4), RegIIIγ*, and *Bdnf* in the ileum (**Fig. 5D)** and increased expression of *Tlr2* in the colon. Therefore, while mucosal barrier function is impaired in both the ileum and colon, evidence of mild immune activation was only observed in the ileum.

**Figure 5.**
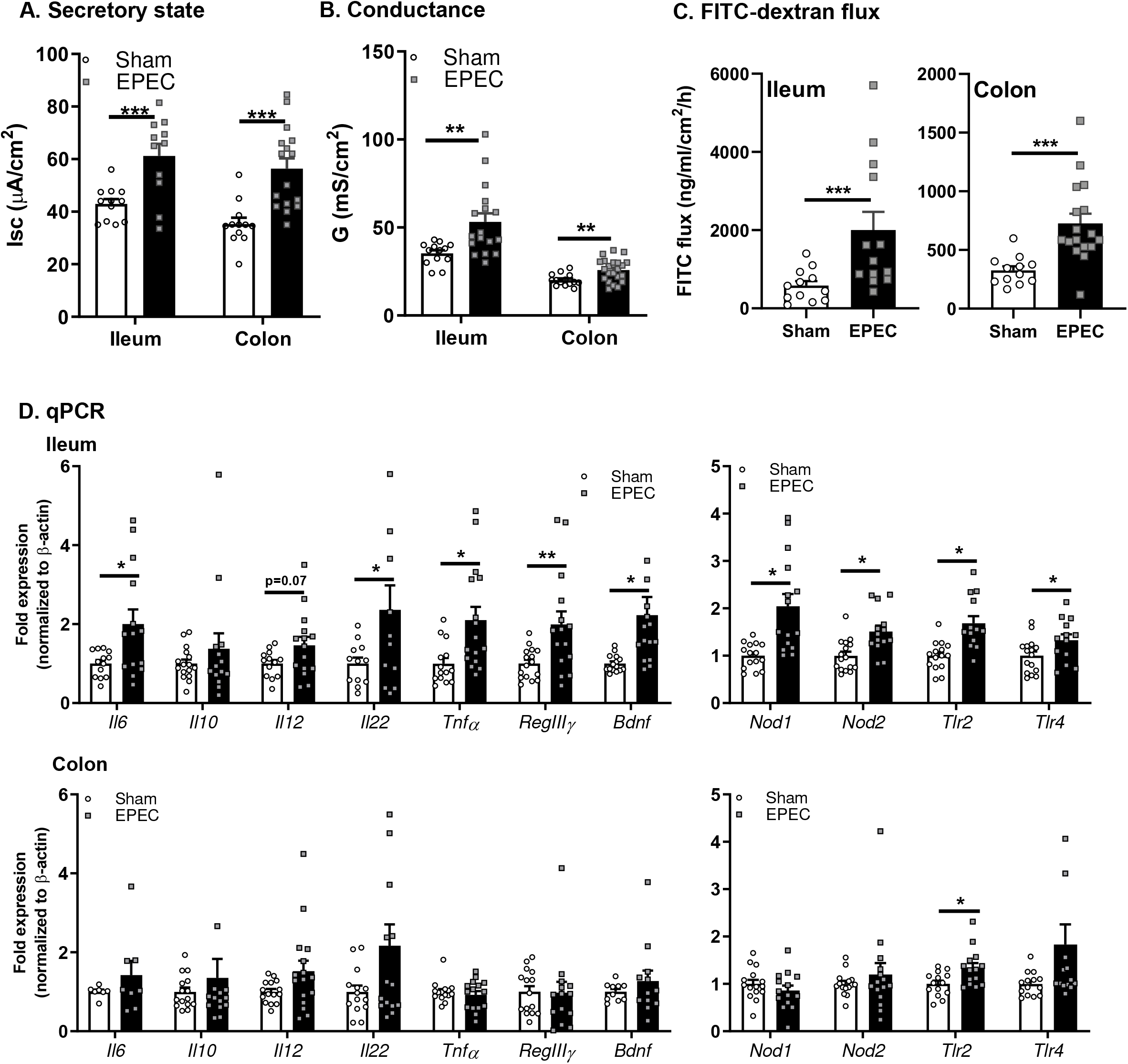
Neonatal EPEC infection caused impaired intestinal physiology in adulthood. Ussing chambers were used to assess (**A**) secretory state (short circuit current [Isc]) and (**B**) tight junction permeability via conductance (G) and (**C**) FITC-dextran flux in both ileum and colon. (**D**) Relative mRNA expression in ileum and colon. N=12-16; *p<0.05, **p<0.01, ***p<0.001 by Student’s T-test, Mann-Whitney and Welch’s correction where appropriate).

### Neonatal EPEC infection causes long-lasting changes to the gut microbiota

In order to characterize changes to the gut microbiota following neonatal EPEC infection, 16S rRNA sequencing of fecal samples from 6-8wk old mice was performed. The Shannon and Chao1 indices suggest a trend towards decreased alpha diversity in neonatally EPEC-infected mice compared to sham-infected controls (Shannon: p=0.09; Chao1 p=0.07; **Fig. 6A**). Using genus level abundance, PLS-DA showed adult sham- and EPEC-infected mice formed separate and distinct clusters (**Fig. 6B**). Phylum level abundances show marked increases in *Actinobacteria* and decreased *Firmicutes* following neonatal EPEC infection compared to sham-infected controls (**Fig. 6C**). At the Family level, several members of the *Lachnospiraceae* and *Ruminococceae* families were decreased in EPEC-infected mice (**Fig. 6D**), whereas *Lactobacillaceae* and *Bifidobacteriaceae* were significantly increased (**Fig. 6D**). Taken together, these findings suggest that neonatal EPEC infection impairs colonization of the gut microbiota, leading to persistent dysbiosis in adulthood.

**Figure 6.**
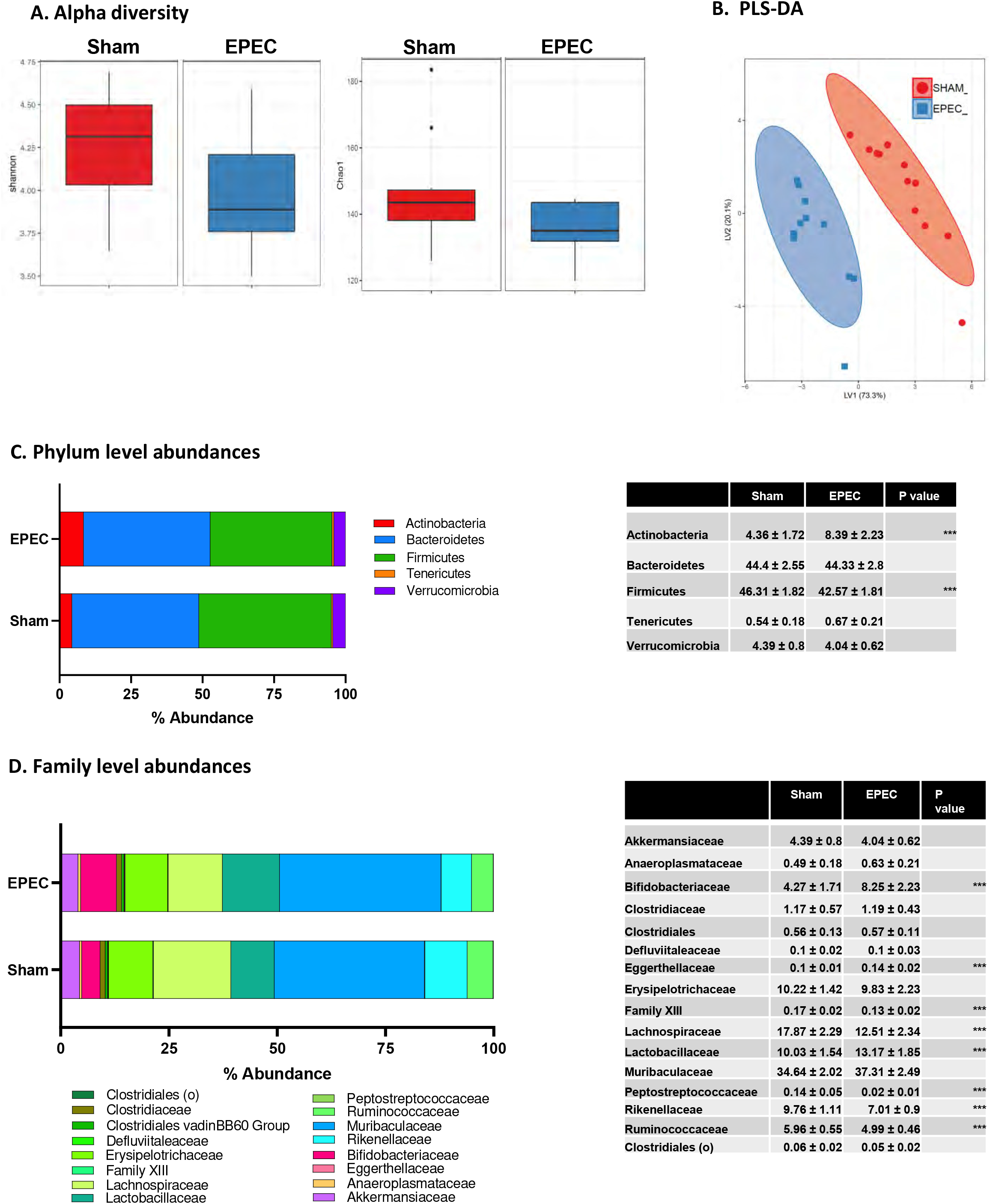
Neonatal EPEC infection causes dysbiosis of the microbiota in adulthood. 16S rRNA Illumina sequencing was performed on fecal pellets to characterize the microbiota. (**A**) Shannon and Chao1 measures of alpha diversity. (**B**) Multivariate analysis using Partial Least Squares Discriminant analysis (PLS-DA) determined using genus level abundances (sham = red; EPEC = blue). (**C**) Phylum level and (**D**) family level abundance, with the highest abundant species presented and statistics outlined in the table. (N=12; ***P<0.001; Holm-Sidak).

## DISCUSSION

Given that establishment of the MGB axis occurs in early life, neonatal enteric infections can potentially have long-lasting impacts on microbiota composition, intestinal physiology, brain, and behavior. Here we demonstrate for the first time, that neonatal EPEC infection can have lasting adverse effects on the MGB axis in mice, well past resolution of the infection by the host. Adult mice neonatally infected with EPEC exhibited cognitive deficits, increased neurogenesis, neuroinflammation, intestinal pathophysiology, and altered gut microbiota composition compared to sham-infected controls. Our findings have implications for better understanding the long-term consequences of enteric infections in early life and other diarrheal diseases in humans.

Multiple studies in adult germ-free mice (12, 35, 36), or following antibiotic administration (37, 38), or bacterial infection (12, 39), have established that changes to the gut microbiota significantly affect the brain and behavior. In developmental studies, maternal antibiotic administration resulted in dysbiosis, coupled with anxiety-like behavior and social behavioral deficits in the pups (11). Similarly, early life stress induced by maternal separation leads to intestinal dysbiosis and pathophysiology (40), as well as behavioral deficits including despair and anxiety-like behavior (41). Here, we found that neonatal infection with a bacterial pathogen led to cognitive deficits without causing anxiety-like behavior in adulthood. These EPEC-induced cognitive deficits required bacterial virulence factors as mice infected with the avirulent ΔescV EPEC strain did not induce memory deficits. These findings indicate that altered host behaviors are due to active infection, and not simply due to Enterobacteriaceae colonization. Additionally, these findings suggest that impairments in host-microbe interactions caused by a bacterial pathogen in early life can disrupt establishment of cognitive functions in adulthood.

Neurogenesis is important for maintaining hippocampal-dependent cognitive function, with perturbations seen in neurodegenerative disorders. Early life experiences, particularly exposure to stress, cause decreased cell proliferation and immature neuron production in adult rats (42). GF mice have increased hippocampal neurogenesis in adulthood, suggesting a role for the microbiota in regulation of this process (43). In our studies, neonatal EPEC infection increased hippocampal BDNF expression and the number of new and immature neurons, suggesting an increase in neurogenesis. Given the importance of neurogenesis in memory formation and retention, increased hippocampal neurogenesis could in part explain the recognition memory deficits observed in neonatally EPEC-infected adult mice.

Neuroinflammation occurs in numerous behavioral and cognitive disorders, with activation of microglia implicated in onset and progression of several neurodevelopmental and neurodegenerative diseases (33). These resident immune cells of the brain survey their microenvironment and are essential for phagocytosis of dying cells; additionally, they play an important role in regulation of neurogenesis and can induce synaptic remodeling (33). Systemic injection of non-pathogenic *E. coli* in neonatal rats caused microglial activation, impaired neurogenesis, and behavior deficits following subsequent immune challenge with lipopolysaccharide in adulthood (44). Similarly, we identified adult memory deficits in neonatally EPEC-infected mice that coincide with long-term neuroinflammation, characterized by immune activation and microglia activation. Taken together, these findings reinforce the susceptibility of the developing neonatal brain to MGB axis defects that persist until adulthood.

Enteric bacterial pathogens are a major cause of diarrhea in humans, resulting in disease in part due to increased ion transport, driving water efflux to flush away the pathogen from the epithelium. Chronic diarrheal illness is associated with a combination of blunted villi, intestinal barrier dysfunction, and submucosal inflammation (45). In addition, early life enteric infections that impair intestinal barrier function have been proposed to cause a shift in the gut microbiota, promoting chronic systemic inflammation and consequently impaired cognitive development (3). In our studies neonatal EPEC infection resulted in intestinal pathophysiology, with increased ileal and colonic ion secretion persisting until adulthood. This pro-secretory state was associated with increased permeability, suggesting increased potential for translocation of bacteria and bacterial products. Indeed, gene expression studies to assess inflammation identified increases in many immune related targets in the ileum of EPEC-infected mice, including *Il6, Tnfα, RegIIIγ, Nod1, Nod2, Tlr2*, and *Tlr4*, which suggests a chronic, low-grade ileal inflammation. These results are keeping with findings in children hospitalized with non-critical infections that have increased intestinal permeability and altered innate immune responses, potentially increasing their risk of subsequent infections (46). This suggests that neonatal bacterial infection can impair establishment of intestinal mucosal barrier function, leading to a chronic mild inflammatory state.

Colonization of the GI tract by the microbiota is crucial for establishment of the intestinal epithelial barrier, maintenance of the mucosal immune response, and resistance to pathogens. The dynamic composition of the microbiota in early life makes it more susceptible to dysbiosis. Here we found that neonatal infection with EPEC decreased the abundance of Firmicutes, particularly the *Lachnospiraceae* family, in adult mice. Given the role of *Lachnospiraceae* in short chain fatty acid production, including butyrate and acetate, decreases in colonization may detrimentally impact host-microbe interactions. Decreased *Lachnospiraceae* was also found in mice exposed to early life stress that exhibited impaired social behavior (47), suggesting these bacteria aid in maintaining gut-brain communication. Taken together, these findings highlight the critical role of neonatal colonization in maintaining host-microbe interactions and suggest that restoration of *Lachnospiraceae* levels may be beneficial for gut-brain axis deficits.

In conclusion, neonatal EPEC infection causes disruption of the MGB axis, characterized by cognitive deficits, neuroinflammation, intestinal pathophysiology, and gut dysbiosis in adulthood. Given the correlation between enteric bacterial infection in children and cognitive function in adulthood, these studies highlight the impact enteric infections can have on the MGB axis and the importance in reestablishing signaling in early life to prevent persistent impairments in adulthood. Future studies assessing the role of beneficial bacteria in ameliorating enteric bacterial infection induced MGB axis deficits in neonates are therefore highly warranted.

## ACKNOWLEDGMENTS

This research was supported by the NIH 1R01AT009365-01 (MGG), 1R21MH108154-01 (MGG), NIH U24-DK092993 (UC Davis Mouse Metabolic Phenotyping Center (RRID:SCR_015361)), a UC Davis MIND Institute Pilot grant NIMH P50HD103526 (MGG), and a training grant NIH T32AI060555 (CH).

## SUPPLEMENTAL FIGURE LEGENDS

**Supplementary Figure 1.**
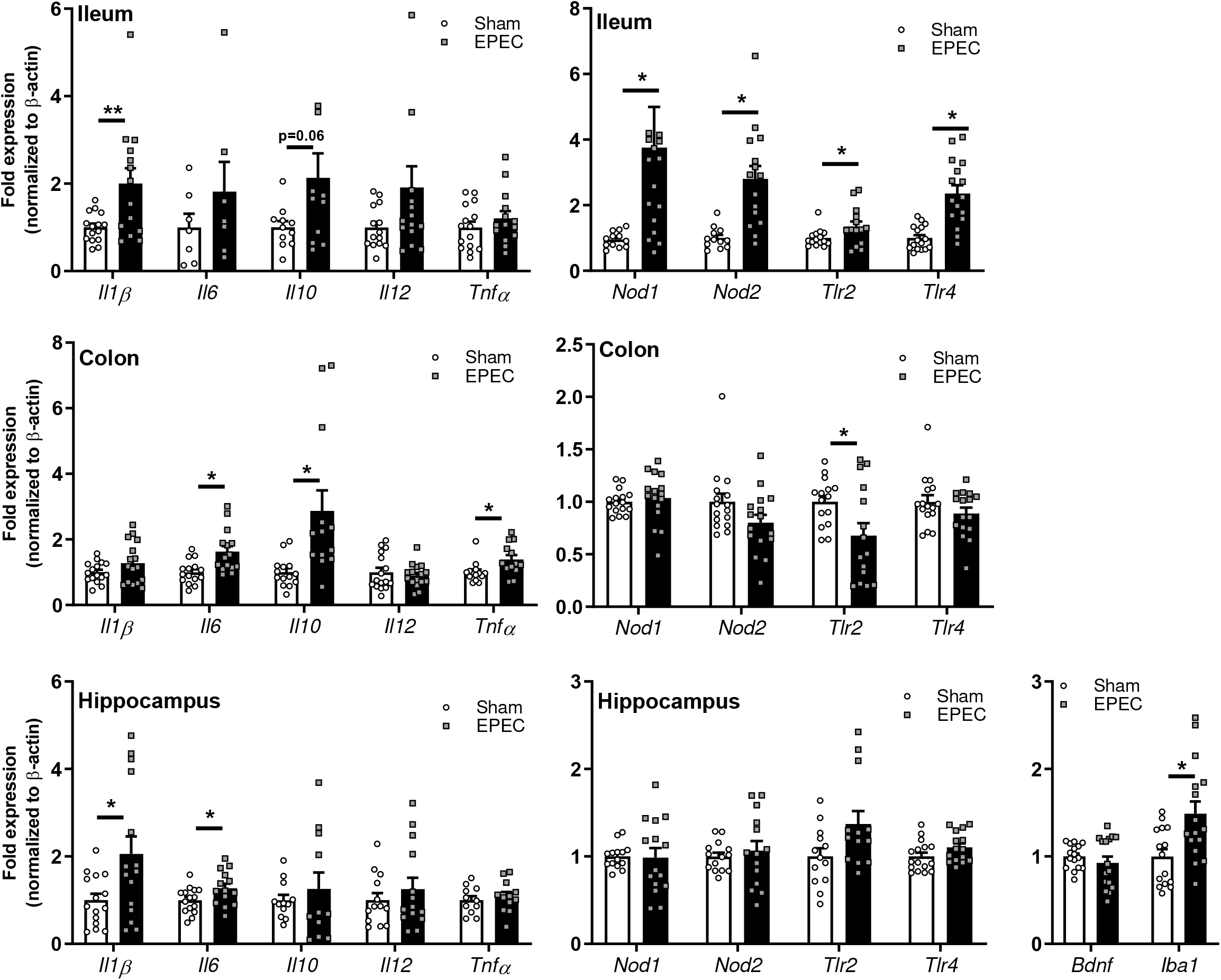
Limited evidence of immune activation at P21 in EPEC-infected mice. Relative mRNA expression in ileum, colon, and hippocampus at P21. (N=12-16; *p<0.05 by Student’s T test (± Welch’s correction), or Mann-Whitney where appropriate).

**Supplementary Figure 2.**
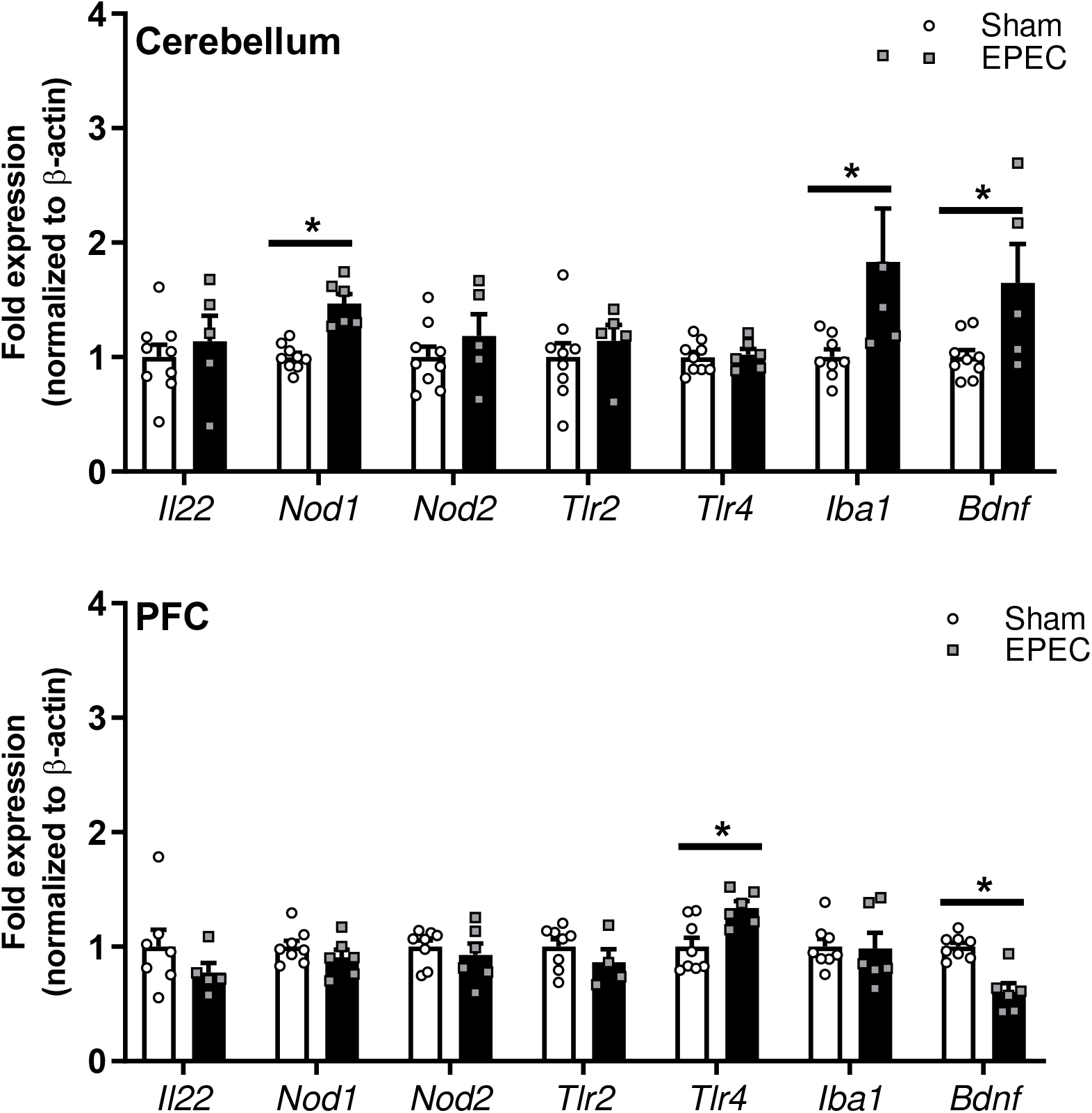
Limited evidence of immune activation in the cerebellum and pre-frontal cortex region in EPEC-infected mice in adulthood. Relative mRNA expression in cerebellum and pre-frontal cortex in adult mice. (N=4-8; *p<0.05 by Student’s T test (± Welch’s correction), or Mann-Whitney where appropriate).

**Supplementary Table 1.**
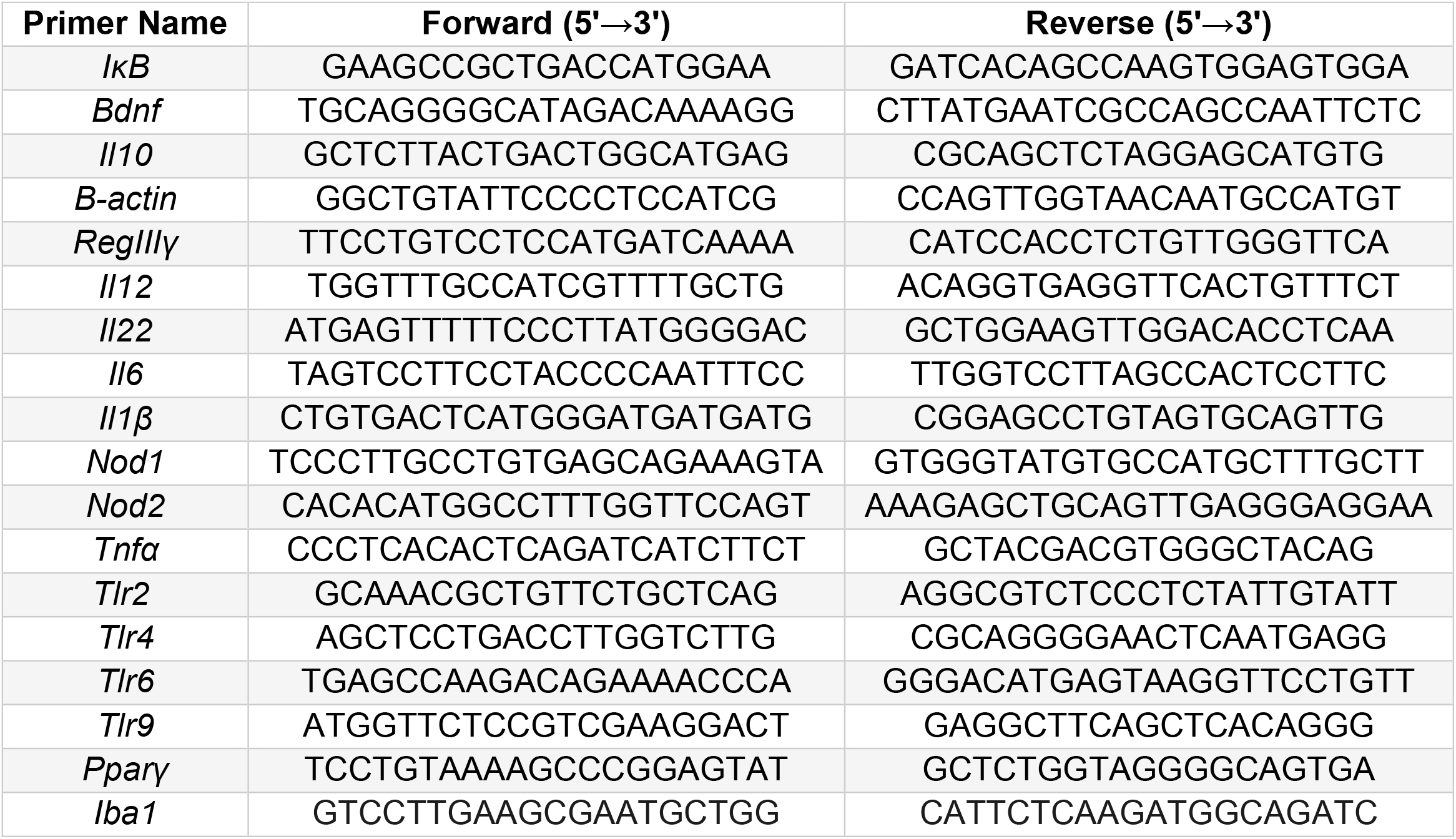
PCR primer sequences

